# Modulation of actin polymerization affects nucleocytoplasmic transport in multiple forms of Amyotrophic Lateral Sclerosis

**DOI:** 10.1101/415901

**Authors:** Anthony Giampetruzzi, Eric W. Danielson, Maryangel Jeon, Valentia Gumina, Sivakumar Boopathy, Robert H. Brown, John E. Landers, Claudia Fallini

## Abstract

Amyotrophic lateral sclerosis (ALS) is a fatal neurodegenerative disease of unknown etiology. Although defects in nucleocytoplasmic transport (NCT) may be central to the pathogenesis of ALS and other neurodegenerative diseases, the molecular mechanisms modulating the nuclear pore function are still largely unknown. Here we show that genetic and pharmacological modulation of actin polymerization disrupts nuclear pore integrity, nuclear import, and downstream pathways such as mRNA post-transcriptional regulation. Importantly, we demonstrate that modulation of actin homeostasis can rescue nuclear pore instability and dysfunction caused by mutant PFN1 as well as by *C9ORF72* repeat expansions, the most common mutations in ALS patients. Collectively, our data link NCT defects to ALS-associated pathology and propose the regulation of actin homeostasis as a novel therapeutic strategy for ALS and other neurodegenerative diseases.

---

Amyotrophic lateral sclerosis (ALS) is a neurodegenerative disease of unknown etiology characterized by progressive loss of motor neurons (MNs). Most ALS cases are sporadic and ∼10% are familial, yet the two classes are clinically indistinguishable suggesting that similar pathways may be responsible for the MN degeneration. Defects in nucleocytoplasmic transport (NCT) have been observed in both cellular and *in vivo* models of ALS and reinforced by pathological evidence in familial and sporadic ALS patients ^1–6^. Nuclear deficiency of mRNA-binding proteins (RBPs) such as TDP-43 and FUS is a pathological hallmark of the disease ^7^, strongly supporting a link between NCT and ALS pathogenesis.

NCT is a tightly regulated process that actively controls the separation between cytoplasmic and nucleoplasmic proteins and RNAs. It is centered on the function of the nuclear pore complex (NPC), a multiprotein complex spanning the whole nuclear envelope and comprised of about 30 different nucleoporins ^8^. Other key players controlling NCT are the small GTPase Ran, its GTPase-activating protein RanGAP1, and the carrier proteins importins and exportins. The cellular distribution of these factors confers directionality to the transport ^9^, while the structural integrity of the NPC and their density across the nuclear envelope modulate the efficiency of the NCT. Interestingly, nucleoporins are the longest lived proteins in the cell, and they are not replaced once the NPC is formed in postmitotic neurons ^10,11^.

Toxic insults such as oxidative stress and protein aggregation have been shown to negatively impact NCT ^12,13^. Many mutant ALS-linked proteins show an increased tendency to aggregate, including SOD1, TDP-43, FUS, and Profilin1 (PFN1). PFN1 is a small actin-binding protein that positively regulates actin polymerization in a formin-dependent manner. ALS-linked mutations in PFN1 render the protein unstable and aggregation prone, leading to the formation of cytoplasmic inclusions ^14,15^. The ability of mutant PFN1 to associate with filamentous (F)-actin is also impaired, and mutant PFN1 MNs show morphological abnormalities, such as smaller growth cones and shorter axons ^15^. However, the molecular mechanisms leading to MN degeneration due to mutations in PFN1 are still unknown. Here we demonstrate that alterations to the actin cytoskeleton caused by mutant PFN1 disrupt the NCT and consequently the normal function of ALS-relevant RBPs, leading to MN dysfunction. Interestingly, modulating actin homeostasis was able to rescue the NCT defects caused by not only mutant PFN1 but also *C9ORF72* repeat expansion, suggesting this could represent a novel therapeutic strategy for ALS.

## RESULTS

### Mutations in PFN1 impair nucleocytoplasmic transport

To investigate whether mutant PFN1 toxicity is associated with nucleocytoplasmic transport (NCT) defects, we examined its effects on the distribution of essential factors controlling this process. To visualize the localization and composition of the nuclear pore complex (NPC) along the nuclear envelope (NE), we stained primary motor neurons (MNs) expressing WT or mutant PFN1 (C71G and G118V) with antibodies recognizing nucleoporins of the FG-Nup family, Nup358/RanBP2, and the transmembrane Nup POM121, given their essential role in regulating NPC structure and function ^16–18^. In WT cells a strong, punctate staining was observed around the nucleus similar to untransfected controls, while a significantly higher percentage of mutant PFN1 MNs showed reduced or absent staining at the NE (Figure 1A-B, S1-S2). Consistent with its known association to the NPC via RanBP2, RanGAP1 localized along the NE in WT cells, while its staining pattern was partially or completely disrupted in mutant PFN1 MNs (Figure 1C). The presence of mutant PFN1 led the transport factor Ran to be abnormally redistributed to the cytoplasm, in contrast to its mostly nuclear localization in WT cells (Figure 1D). This effect was more pronounced in cells containing visible inclusions, although MNs with no obvious aggregates still had Ran C:N ratios significantly higher than WT values. No co-aggregation of any of the tested proteins with mutant PFN1 was observed (Figure 1E-G), and no changes in RanGAP1 SUMOylation, which is necessary for its association with the NPC ^19^, were detected (Figure 1H). Similarly, no difference in the overall levels of the tested nucleoporins was observed in all conditions, while a slight reduction in Ran levels was present in PFN1^C71G^ MNs (Figure S3). We did not observe changes to the localization of karyopherins, with the exception of a small reduction in Exportin 1 (XPO1) levels (Figure S4).

**Figure 1.**
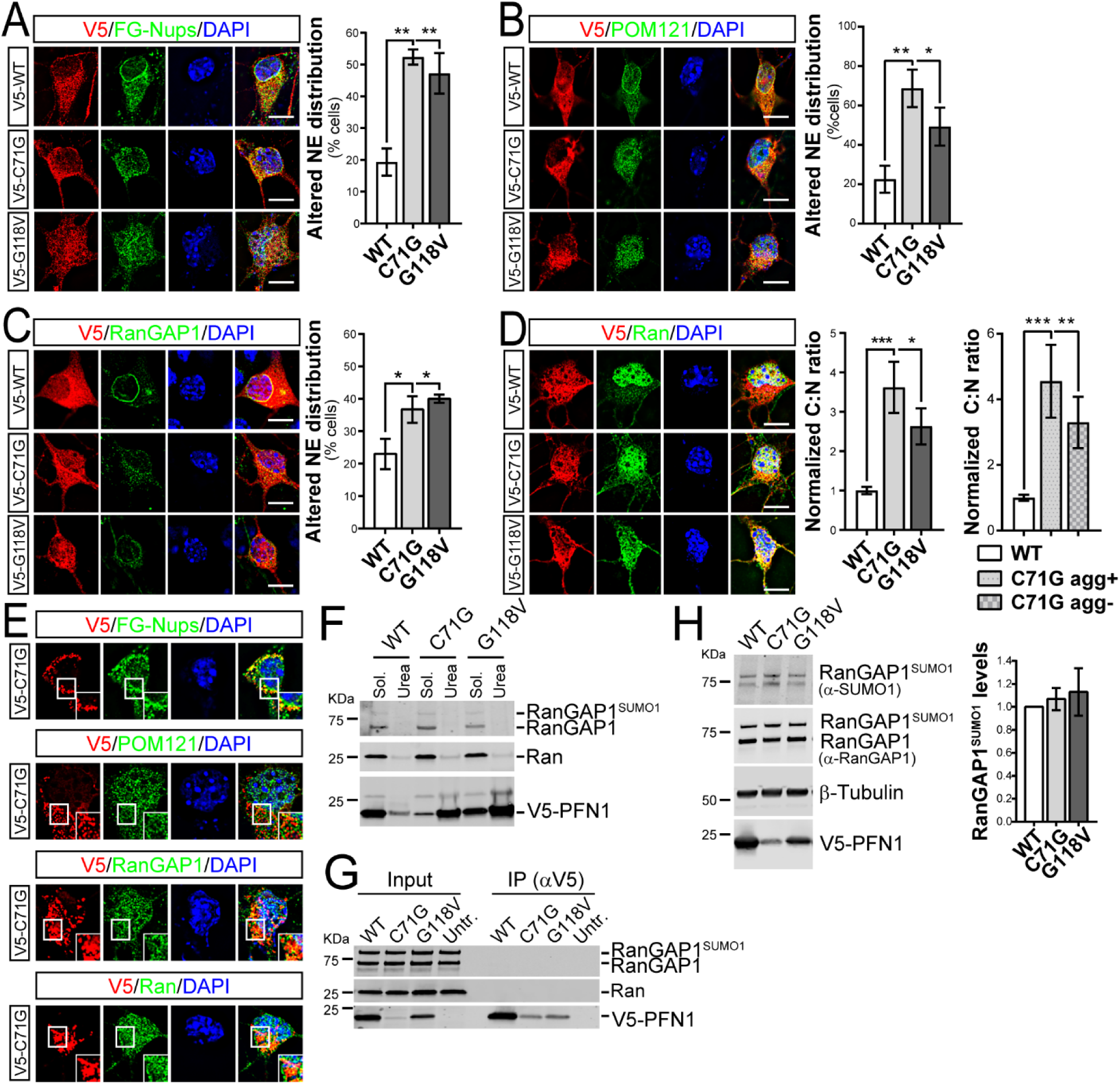
Mutant PFN1 alters the composition and density of NPCs. **A-C.** FG-NupsPOM121 (*B, green*), and RanGAP1 (*C, green*) localization to the NE is altered in a higher percentage of MNs expressing V5-tagged mutant PFN1 vs WT control (*red*). **D.** Ran (*green*) cytoplasm to nucleus (C:N) ratio is increased in MNs expressing V5-WT or mutant PFN1 (*red*), regardless of the presence of aggregates (agg), indicating possible functional defects in the segregation of cytoplasmic and nuclear proteins. **E.** The presence of PFN1^C71G^ aggregates (*red*) in MNs does not lead to co-aggregation of FG-Nups, POM121, RanGAP1, or Ran (*green*). **F**. No difference in the solubility of Ran (*middle panel*) or RanGAP1 (*top panel*) caused by the expression of PFN1 mutants when assayed in HEK293 cells using detergent-based cellular fractionation. Triton X-100 (2%) and urea (8M) were used to extract the soluble and insoluble fraction, respectively. **G.** Representative blot of a co-immunoprecipitation (co-IP) assay between V5-tagged WT or mutant PFN1 and RanGAP1 (*top panel*) or Ran (*middle panel*). No bands were detected in the IP pellet, suggesting lack of interaction. **H.** Representative western blot and quantification showing unchanged levels of SUMO1-modified RanGAP1 in the presence of mutant PFN1. Both antibodies against SUMO1 (*top panel*) and RanGAP1 (*second panel*) detect a band ∼80KDa corresponding to SUMOylated RanGAP1. V5 antibody (*bottom panel*) shows the expression of the V5-tagged PFN1 protein, while β-tubulin was used as a loading control. Bars are mean±SEM; \**p*<0.05, \*\**p*<0.01, \*\*\**p*<0.001. DAPI (*blue*) was used to detect the nucleus and assess cell health. Scale bars: 10µm. See also Figure S1-S4, and Table S1 for details on statistics.

### Mutant PFN1 alters the structure of the nuclear membrane

Since the NPC is tightly connected with the nucleoskeleton, we further investigated the effect of mutant PFN1 on the nuclear structure using transmitted electron microscopy. Similarly to what was observed in cells expressing TDP-43 C-terminal fragment ^1^, we found that the expression of either V5-tagged or GFP-tagged PFN1^C71G^ in Neuro2a cells led to severe defects in the structure of the nucleus, with the presence of frequent folds, invaginations, and protrusions that were never observed in untransfected or PFN1^WT^-transfected cells (Figure 2A). This observation was confirmed by immunofluorescence analysis of Lamin A/C, which showed non-uniform and irregular staining in cells expressing mutant PFN1 (Figure 2B), and a trend toward reduced levels in cells with abnormal distribution.

**Figure 2.**
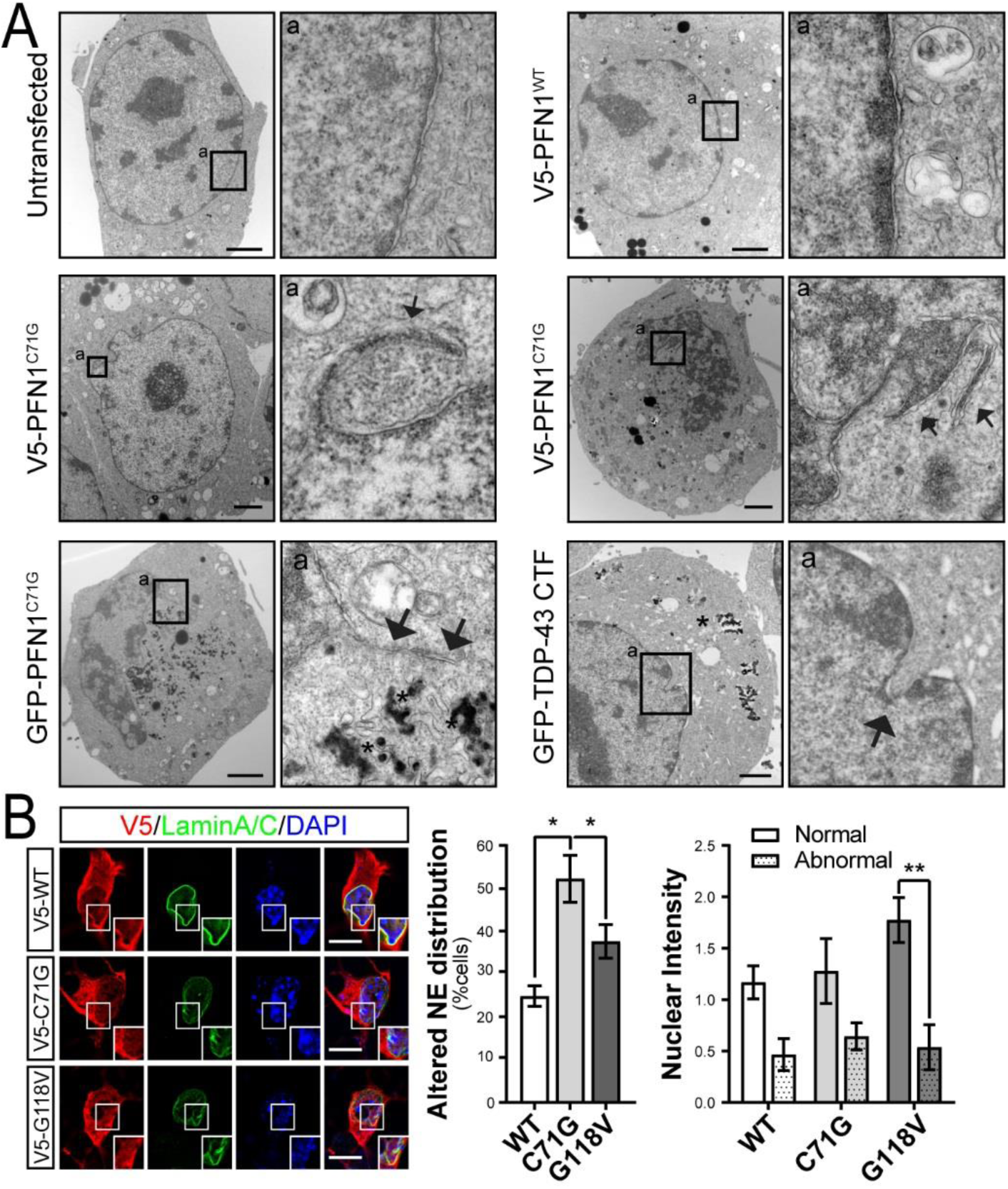
The nuclear membrane integrity is compromised by mutant PFN1. **A.** Transmission electron microscopy shows the presence of protrusions and folds (arrows) in Neuro2a cells expressing V5- or GFP-tagged PFN1^C71G^ compared to untransfected (Untr.) or PFN1^WT^ cells, similar to what observed in the presence of TDP-43 C-terminal fragment (CTF). Arrows point to anomalous membrane structures in the nucleus. Aggregates (asterisks) are visible as dark amorphous structures in the cytoplasm. **B.** Lamin A/C (green) distribution at the NE is altered in a higher percentage of mutant PFN1 MNs. Overall levels of Lamin A/C also appear slightly reduced in cells with abnormal staining. Boxes indicate areas enlarged in insets. Bars are mean±SEM; \**p*<0.05, \*\**p*<0.01. DAPI (blue) was used to detect the nucleus and assess cell health. Scale bars: 2µm in A, 10µm in B. See also Figure S1 and S3, and Table S1 for details on statistics.

### Nuclear import is greatly reduced by mutant PFN1

To further explore the functional consequences of the structural defects caused by mutant PFN1 on the NCT, we measured the rate of nuclear import by live cell imaging using a NLS-NES-mCherry (Shuttling (S)-mCherry) reporter ^20^. Cortical neurons were co-transfected with GFP or GFP-tagged PFN1 and S-mCherry. S-mCherry localized mainly to the cytoplasm in all conditions due to the stronger effect of the nuclear export signal (NES) compared to the nuclear localization signal (NLS)(Figure 3E).Thirty-six hours after transfection, cells were treated with Leptomycin B (LMB) – a selective inhibitor of Exportin 1 – to inhibit nuclear export, leading to a measurable time-dependent accumulation of the reporter in the nucleus. We found that the expression of mutant PFN1 led to a significant reduction in import rates compared to both GFP- and PFN1^WT^-transfected cells (Figure 3A-C), and an increase in the percentage of non-responder cells (39% in C71G vs 6% WT) (Figure 3D). In all, these data suggest that mutant PFN1 may affect nuclear stability, altering either the composition and/or the number of functional NPCs, and thus leading to severe nuclear import defects.

**Figure 3.**
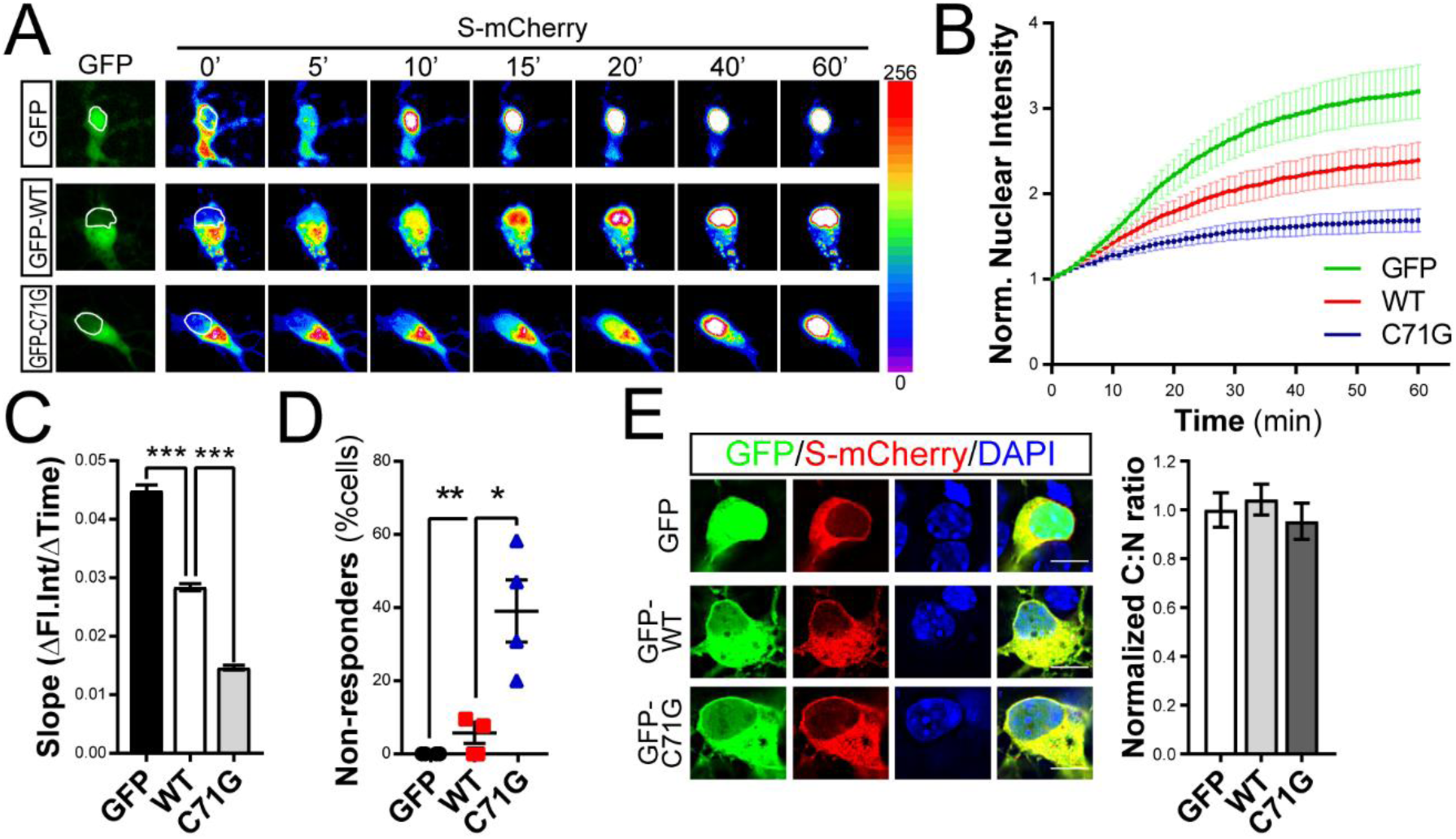
Mutant PFN1 alters the efficiency of nuclear import. **A-B.** Time-lapse images and quantification of S-mCherry import dynamics in cortical neurons expressing GFP or GFP-tagged PFN1 (*green*) and treated with LMB. S-mCherry levels are shown as 16-bit heat map. Dashed lines indicate the nucleus. **C-D.** Regression analysis of S-mCherry kinetics (*C*) and percentage of cells whose nuclear levels did not rise above 1.2 folds over initial values upon the application of LMB (i.e. non-responders, *D*). **E.** S-mCherry (*red*) C:N ratio is not affected by mutant PFN1 (*green*) in untreated cells. GFP alone was used as control, and localizes to both nucleus and cytoplasm. Bars are mean±SEM; \**p*<0.05, \*\**p*<0.01, \*\*\**p*<0.001. DAPI (*blue*) was used to detect the nucleus and assess cell health. Scale bars: 10µm. See Table S1 for details on statistics.

### mRNA post-transcriptional regulation is impaired in mutant PFN1 cells

One of the main classes of proteins that shuttle between the nucleus and the cytoplasm are RBPs which control the post-transcriptional fate of mRNAs. Thus, we investigated the impact of mutant PFN1-dependent disturbance to NCT on the distribution of RBPs by quantifying the nuclear and cytoplasmic levels of the mostly nuclear proteins TDP-43 and FUS, and of the mostly cytoplasmic proteins SMN and FMRP. As expected, the majority of TDP-43 and FUS was nuclear in WT-expressing MNs, while mutant PFN1 expression led to a shift in the C:N ratio of both proteins (Figure 4A-B). Again, no effect of PFN1^WT^ expression was observed on the localization of the proteins compared to untransfected controls (Figure S5). On the contrary, there we no quantifiable changes to the distribution of the mostly cytoplasmic FMRP and SMN proteins, although we observed fewer SMN-positive nuclear gems (Figure S6). We also observed a significant reduction in TDP-43 localization to the proximal motor axon (Figure S5) and an increase in TDP-43 aggregation, possibly mediated by TDP-43’s illegitimate interaction with mutant PFN1 (Figure S7). Together, these data suggest that mutant PFN1 perturbs the distribution of ALS-relevant nuclear RBPs by destabilizing the NPC and impairing nuclear import. To investigate the downstream consequences of PFN1-mediated mislocalization of such RBPs, we focused as a proof of principle on TDP-43’s well-studied regulatory activity on the axonal localization of the *neurofilament L* (*Nefl*) mRNA ^21^ and on the splicing of *POLDIP3* mRNA ^22^. Quantitative fluorescence *in situ* hybridization was performed in MNs expressing either GFP or GFP-tagged WT or C71G PFN1 (Figure 4C, S8). While no difference in the somatic levels of the *Nefl* mRNA was detected, its axonal levels were severely reduced in C71G-expressing MNs. Immortalized lymphoblast cell lines derived from 4 controls and 3 patients carrying *PFN1* mutations ^15^ were used to evaluate the abundance of two alternatively spliced *POLDIP3* variants – S1 and S2 – by RT-PCR (Figure 4D, S8). In all ALS lines, the relative levels of isoform S2 were significantly increased over control cells. Together these data suggest a loss of function for TDP-43 in mutant PFN1 cells, possibly due to its nucleocytoplasmic redistribution.

**Figure 4.**
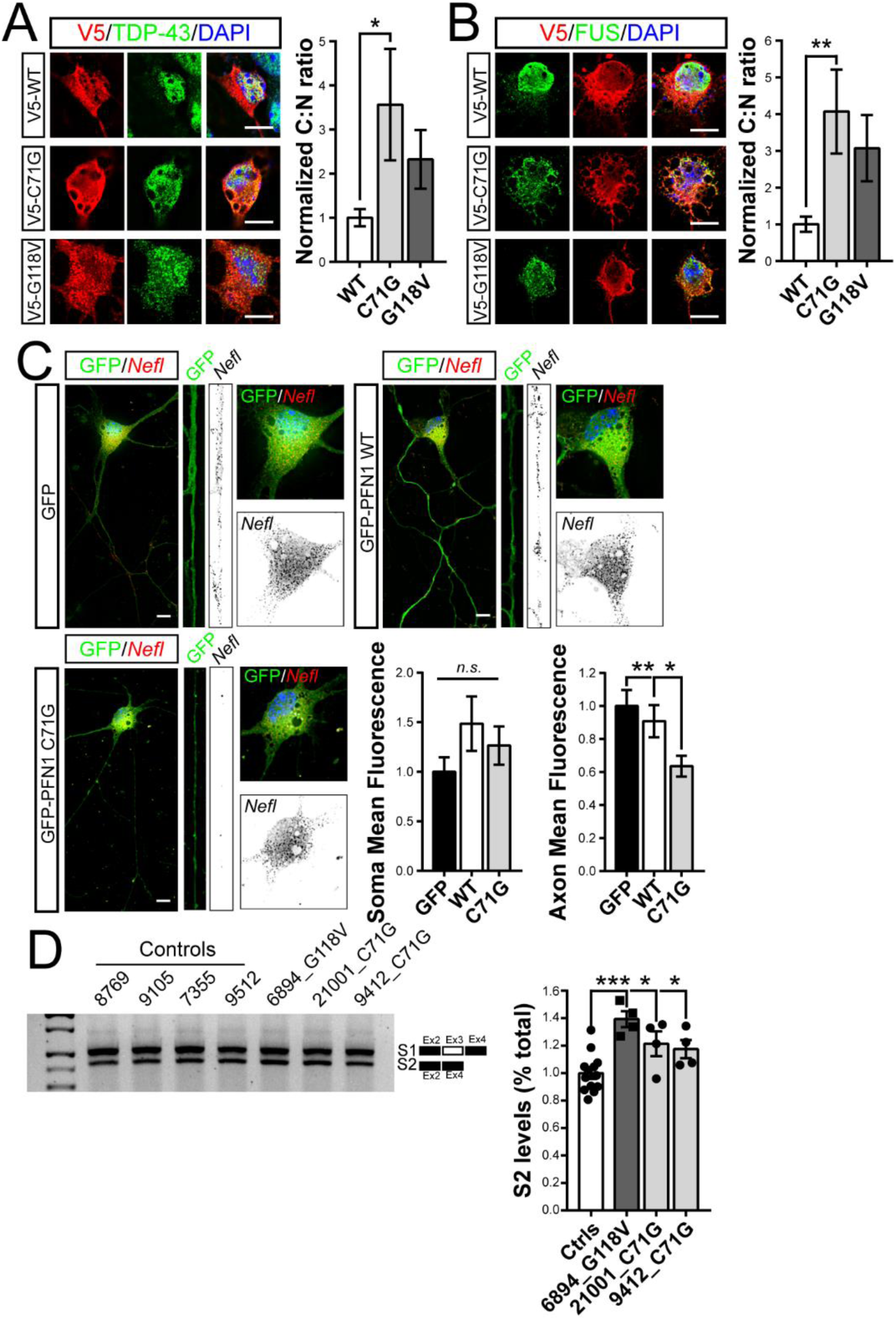
Mutant PFN1 perturbs RBPs cellular distribution and function. **A-B.** Mutant PFN1 (*red*) causes redistribution of nuclear RBPs TDP-43 (*A, green*) and FUS (*B, green*) to the cytoplasm, as quantified by their cytoplasm to nucleus (C:N) ratios. **C.** RNA-FISH analysis reveals that the *Nefl* mRNA (*black*) levels in the axon, but not in the cell soma, are significantly reduced due to the expression of GFP-PFN1^C71G^ (*green*) compared to GFP or GFP-PFN1^WT^-expressing MNs. **D.** Representative DNA gel and quantification of the levels of the *POLDIP3* S2 variant over total *POLDIP3* levels (S1+S2) in control (Ctrl) versus mutant PFN1 lymphoblast lines. Bars are mean±SEM; \**p*<0.05, \*\**p*<0.01, \*\*\**p*<0.001. DAPI (*blue*) was used to detect the nucleus and assess cell health. Scale bars: 10µm. See also Figure S5-S8, and Table S1 for details on statistics.

### Inhibition of nuclear export improves ALS relevant disease phenotypes

To assess the causality between NCT disturbance, RBP mislocalization, and MN pathology, we investigated the potential of nuclear export inhibitor KPT-276, a well-established selective inhibitor of XPO1 ^6,23,24^, to rescue PFN1-dependent defects. First, WT or C71G expressing MNs were treated with 50nM KPT-276 or DMSO 6 hours prior to fixation, and the C:N ratio of TDP-43 was determined. KPT-276 treatment was able to fully rescue TDP-43 cytoplasmic mislocalization (Figure 5A), confirming successful inhibition of nuclear export. Next, we sought to determine the potential of KPT-276 treatment to rescue axonal outgrowth defects previously described in mutant PFN1 MNs ^15^. MNs expressing mutant PFN1 and treated vehicle alone had significantly shorter axons than WT cells, while KPT-276 treatment fully rescued the defect (Figure 5B). Similar rescue was observed by measuring the rate of axon growth by live cell imaging over the course of 1 hour (Figure 5C, Movie S1). Together, these data support a direct link between NCT, mRNA regulation, and PFN1-dependent ALS pathology.

**Figure 5.**
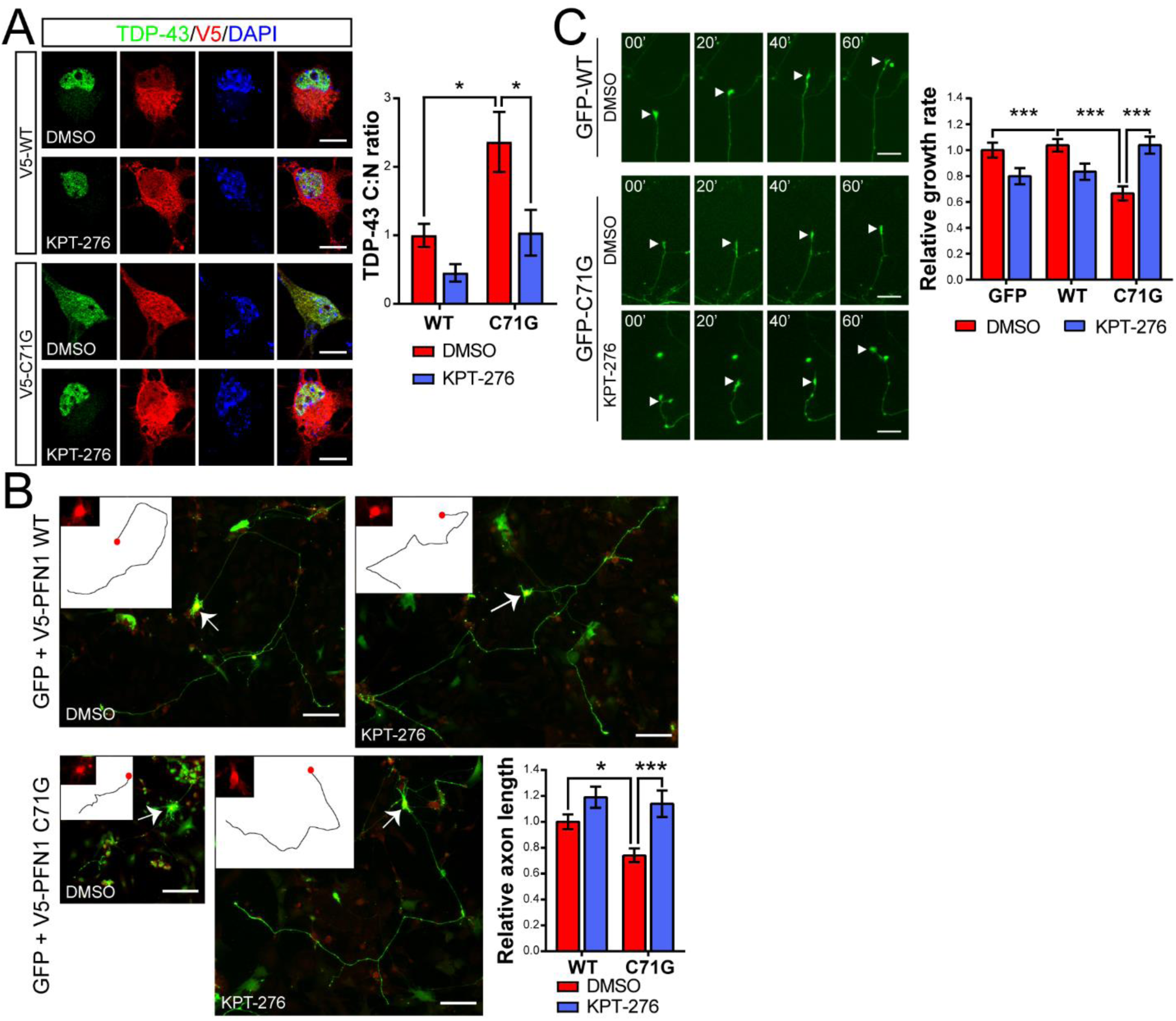
Inhibition of nuclear export rescues ALS-associated and PFN1-dependent pathology. **A.** KPT-276 treatment rescues TDP-43 (*green*) C:N ratio in MNs expressing mutant PFN1 (*red*). **B.** MNs expressing PFN1 C71G had significantly shorter axons compared to WT-expressing cells. This defect was fully rescued by KPT-276 treatment. Insets show the tracing of the primary axon and the expression of V5-PFN1 in the cell body (*red*, arrows). **C.** Representative time lapse images and quantification of axon outgrowth in the presence or absence of KPT-276. A full rescue of the outgrowth defects was observed following KPT-276 treatment. Bars are mean±SEM; \**p*<0.05, \*\*\**p*<0.001. DAPI (*blue*) was used to detect the nucleus and assess cell health. DMSO was used as vehicle control. Scale bars: 10µm. See also Table S1 for details on statistics.

### Modulation of actin polymerization modifies NCT in mutant PFN1 MNs

PFN1’s main cellular function is to promote actin polymerization by facilitating formin-based actin nucleation and elongation, a function impaired by the presence of ALS-linked mutations ^14,15^. Thus, we hypothesized that PFN1-mediated disruption of the actin network could directly interfere with protein nuclear shuttling by affecting nuclear stability. Supporting this hypothesis, we observed a severe mislocalization of both RanGAP1 and Ran in MNs treated with the actin depolymerizing drug Latrunculin A (LatA) (Figure 6A-B and S9), resembling the phenotypes identified in mutant PFN1 MNs. To further this observation, we used a complementary approach and attempted to rescue NPC defects in PFN1^C71G^ MNs by positively modulating actin polymerization. To this end, we took advantage of the ability of formins to promote actin polymerization, albeit at a slower rate, even in the absence of functional PFN1 ^25^. Overexpression of a constitutively active form of the formin mDia1 (FH1-FH2 domains) in mutant PFN1 MNs was indeed able to restore normal actin homeostasis (Figure S9), without changing the aggregation propensity of PFN1^C71G^. Importantly, while mutant PFN1 MNs expressing GFP alone had significantly higher frequency of disrupted RanGAP1 staining, as previously observed (see Figure 1), the expression of GFP-mDia1 fully rescued the defect (Figure 6C). Similarly, GFP-mDia1 expression was able to rescue TDP-43 mislocalization (Figure 6D) in mutant PFN1 MNs. To assess whether actin modulation could also rescue the function of the nuclear pore, we again measured import dynamics of S-mCherry in the presence of mDia1 overexpression. A significant improvement of import dynamics was observed in PFN1^C71G^ neurons following mDia1 expression (Figure 6E-H), suggesting that changes in actin homeostasis can influence the stability of the NPC and/or NE, affecting protein shuttling.

**Figure 6.**
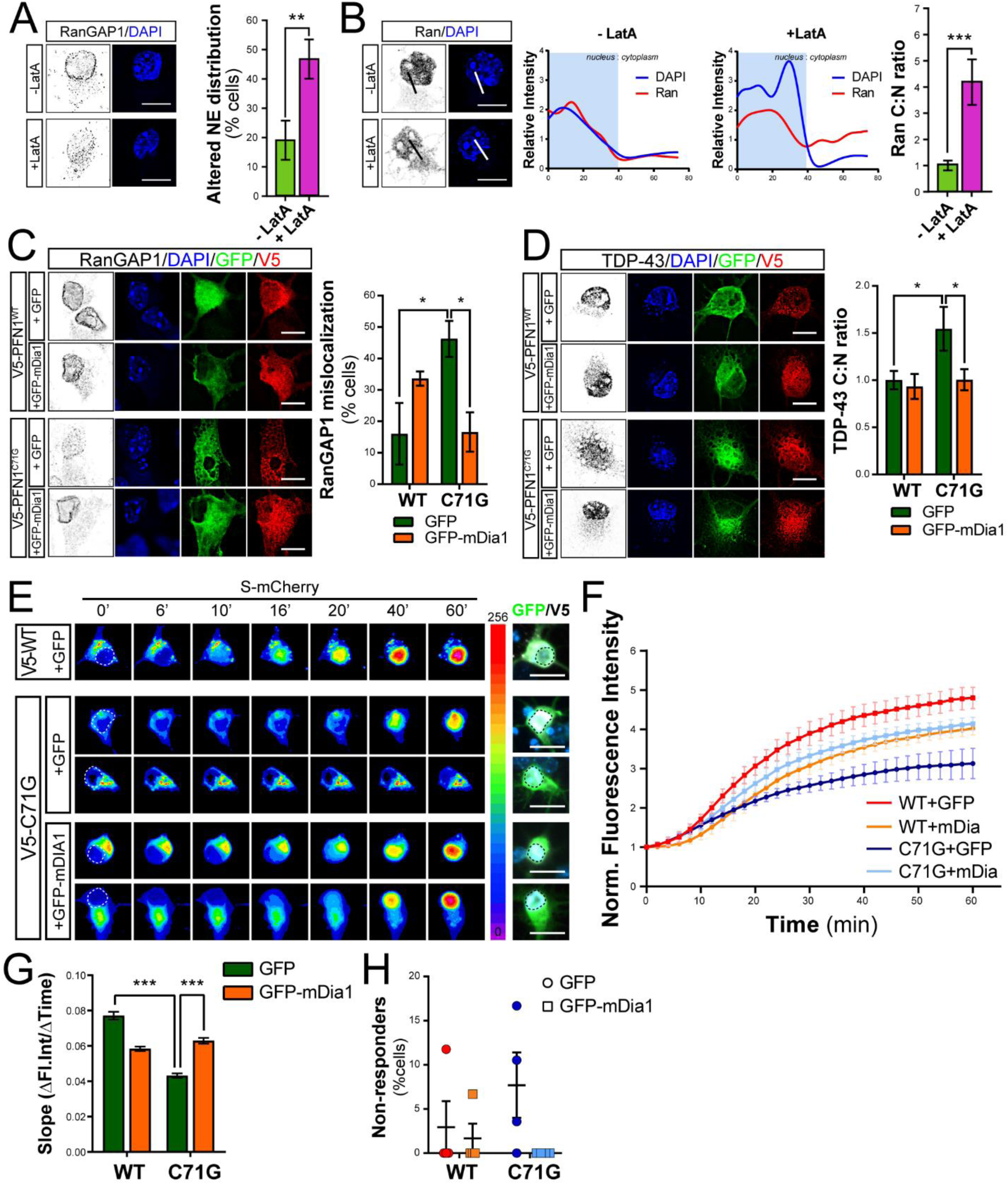
Actin homeostasis is a significant modulator of NPC structure and function. **A-B.** RanGAP1 (*A*) and Ran (*B*) localization is disrupted by Latrunculin A (LatA) treatment. Line plots of Ran and DAPI intensity are shown. **C-D.** Representative images and quantification show rescue of RanGAP1 (*C*) and TDP-43 (*D*) mislocalization due to the overexpression of GFP-mDia1 in MNs expressing PFN1C^71G^. **E-F.** Time-lapse images and quantification of S-mCherry import dynamics upon treatment with LMB in cortical neurons expressing GFP or GFP-mDia1 (*green*) and V5-tagged PFN1 (*white*). S-mCherry levels are shown as 16-bit heat map. Dashed lines indicate the nucleus. Cells were post-fixed and stained to verify expression of V5-PFN1 constructs (*white*). **G-H.** Regression analysis of S-mCherry kinetics (*G*) and percentage of non-responder cells (*H*). Bars are mean±SEM. \**p*<0.05, \*\**p*<0.01,\*\*\**p*<0.001. DAPI (*blue*) was used to detect the nucleus and assess cell health. Scale bars: 10µm. See also Figure S9 and Table S1 for details on statistics.

### Modulation of actin polymerization rescues NCT defects in C9ORF72-ALS

Next, we sought to investigate whether modulating the actin cytoskeleton could be beneficial in other forms of ALS characterized by defects to the nuclear pore function, such as C9ORF72-ALS. Thus, we transfected MNs with a synthetic construct expressing 80 GGGGCC repeats (G4C2)80, that had been previously shown to induce the presence of RNA foci and dipeptide formation ^26^. The expression of this construct did not result in any obvious difference in F-actin levels at the growth cone under our experimental conditions (Figure S10). However, impaired actin dynamics due to decreased functional cofilin were observed in primary MNs and patient derived cells and tissues harboring a G4C2 repeat expansion in the *C9ORF72* locus ^27^. As anticipated, expression of (G4C2)80 repeats in MNs led to the loss of RanGAP1 localization to the NE with no change to its overall levels (Figure 7A and S10) ^6^. Interestingly, overexpression of the constitutively active form of mDia1 fully rescued the phenotype. Notably, mDia1 expression was also able to rescue functional import defects as detected by S-mCherry dynamics (Figure 7B-D). Together, these results suggest that modulation of actin homeostasis directly modifies the stability of the NPC and/or NE, protecting it from disruption caused by ALS-associated gene mutations.

**Figure 7.**
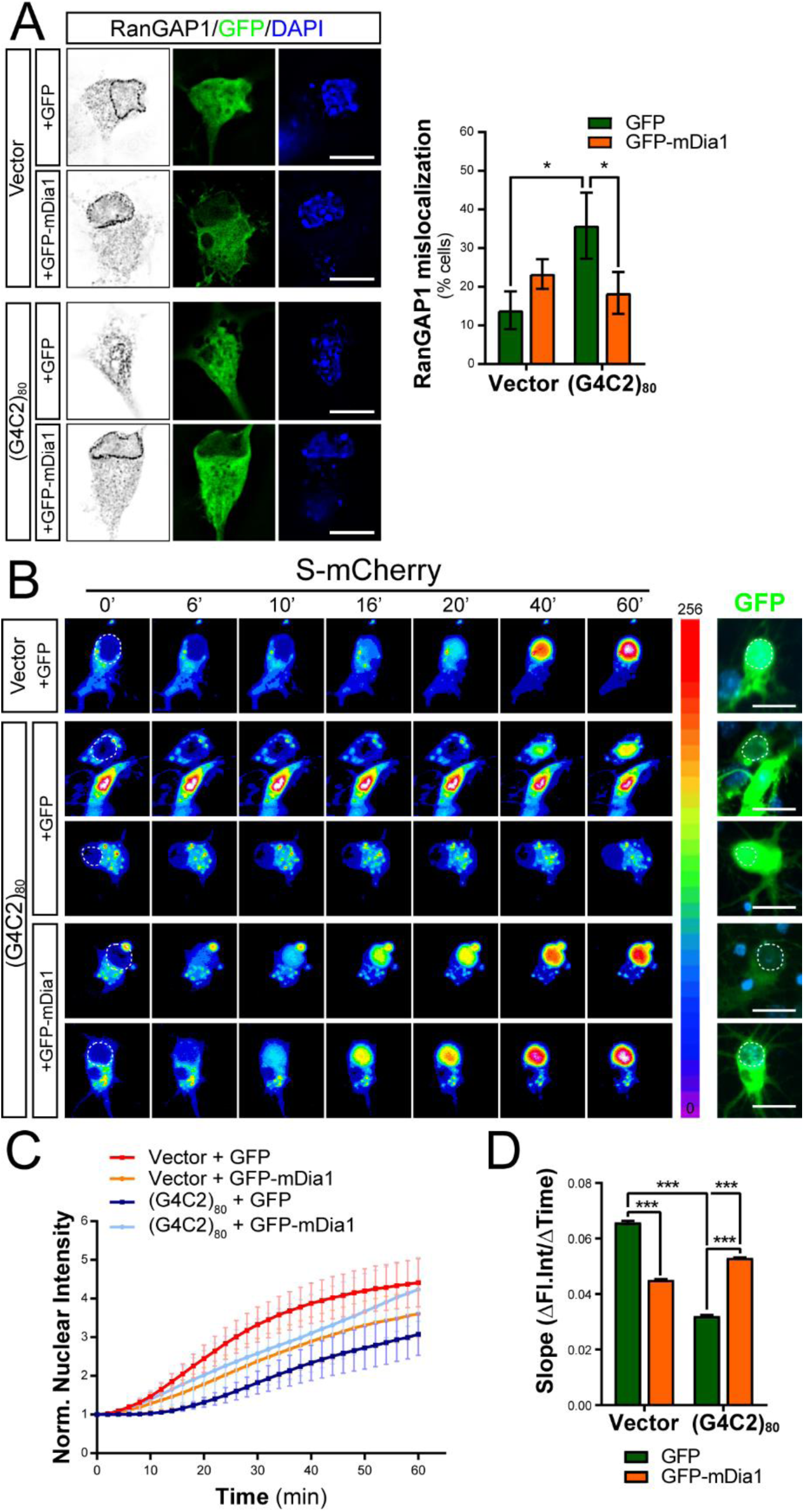
Actin modulates NPC function in C9ORF72-ALS. **A.** Significant rescue of RanGAP1 (*black*) mislocalization to the NE in cells expressing the *C9ORF72* repeat expansion (G4C2)^80^ was achieved due to the overexpression of GFP-mDia1 compared to GFP alone (*green*). **B-D.** Time-lapse images (*B*), quantification (*C*) and regression analysis (*D*) of S-mCherry import dynamics in cortical neurons expressing GFP or GFP-mDia1 (*green*) and (G4C2)^80^. S-mCherry levels are shown as 16-bit heat map. Dashed lines indicate the nucleus. Scale bars: 10µm. Bars are mean±SEM. \**p*<0.05, \*\*\**p*<0.001. DAPI (*blue*) was used to detect the nucleus and assess cell health. See also Figure S10 and Table S1 for details on statistics.

## DISCUSSION

ALS is a heterogeneous disease and mutations in a wide array of genes are associated with ALS pathogenesis ^28^. Although a common mechanism linking these different gene mutations may not be immediately apparent, recent findings from several groups point at defects in NCT as central to ALS pathology ^29^. Here we explore the hypothesis that alterations in cytoskeletal stability caused by ALS-associated mutations in the actin-binding protein PFN1 would similarly impact the function of the NPC.

Our immunofluorescence data in primary MNs show that mutant PFN1 leads to a severe impairment of the NPC structure and/or number, marked by the loss of nucleoporins from the nuclear envelope, as well as of its function, as indicated by the dissolution of the nucleocytoplasmic gradient of Ran, reduced import rates, and cytosolic mislocalization of nuclear RBPs TDP-43 and FUS. No changes in the distribution of cytoplasmic shuttling proteins (i.e. SMN, FMRP, S-mCherry, and PFN1 itself) were detected, suggesting that mutant PFN1 mostly affected nuclear import versus export. To investigate how mutant PFN1 disrupts the NCT, we tested whether it directly affected RanGAP1 protein levels or its association with the nuclear pore, as suggested in flies where loss of function of PFN1 ortholog *Chic* was associated with selective RanGAP1 irregular localization to the NE ^30^. However, we did not detect any change in RanGAP1 levels or SUMOylation, which is necessary for its binding to the NPC. Next, we investigated whether PFN1 aggregation could sequester NCT factors, thus causing impairment in nuclear shuttling as shown for synthetic amyloid aggregates and TDP-43 ^1,12^. Again, we were not able to detect increased insolubility or co-aggregation of the factors tested with PFN1 inclusions. Additionally, while TDP-43 aggregates are cross-seeded by PFN1 ^15,31^, we observed mislocalization of nucleoporins and Ran even in the absence of PFN1 inclusions, suggesting that TDP-43 mislocalization is a consequence rather than the cause of NCT disturbance. Finally, we excluded from our analyses any of the rare cells with spontaneous stress granule-like formation due to PFN1 overexpression ^32^, thus eliminating the possibility that stress granules would induce NCT defects in our experimental system ^13^. A more likely scenario is that mutant PFN1-dependent disruption of the actin cytoskeleton destabilizes the nuclear membrane, leading to NCT defects. This is supported by the observation that actin depolymerization caused by Latrunculin A was sufficient to cause severe changes in the cellular distribution of Ran and RanGAP1, and that the overexpression of the constitutively active form of mDia1, by promoting filamentous actin polymerization and without changing PFN1 aggregation propensity, was able to structurally and functionally rescue the NPC. Intriguingly, we found that mDia1 rescue was not limited to mutant PFN1 cells, but extended to cells expressing the *C9ORF72* repeat expansion, the most common mutation found in ALS patients. Although the mechanisms driving this rescue are not fully understood yet, we speculate that actin may act by modulating the integrity of the NE via the function of the linker of nucleoskeleton and cytoskeleton (LINC) complex ^33^, a multiprotein complex that physically connects lamins with actin and microtubules. Of note, mutations in LINC components cause cerebellar ataxia ^34,35^, suggesting that this complex plays an important role in the maintenance of neuronal function and survival, possibly through the modulation of NPC function. Future work will be required to reveal how changes to actin homeostasis affect the LINC complex and its functional consequences on the NCT and how subtle changes to actin polymerization could be employed as a strategy to modulate NCT function.

It remains an open question whether the NCT defects observed in our model and others are a cause or a mere effect of the degenerative process. Our data strongly argue for the NPC dysfunction being a primary driving force of ALS-associated cell toxicity. Indeed, we found that pharmacological inhibition of nuclear export rescued PFN1-dependent cellular pathology, such as stunted axon growth. Together with previous observation connecting deficiencies in NCT to axonal guidance defects in flies ^36^, our data are particularly interesting as they functionally link the cytoskeleton with NCT and mRNA post-transcriptional regulation. Axon growth is a complex process that requires active cytoskeletal growth ^37^, fueled by the increased expression, transport, and local translation of several cytoskeletal-related mRNAs ^38,39^. Defects in the localization and local translation of mRNAs at the tip of axons and dendrites has been described in other neurological diseases, such as SMA and Fragile X syndrome ^40,41^, supporting the relevance of these defects in disease pathogenesis. Our data suggest that PFN1-dependent impairment of NCT and consequent mislocalization/loss of function of nuclear RBPs leads to impaired mRNA regulation, of which *POLDIP3* mRNA splicing and *Nefl* mRNA axonal localization are two examples. Together, these data support our hypothesis that PFN1-mediated and actin-dependent disruption of NCT may have severe downstream consequences on general mRNA regulation, leading to a vicious cycle that eventually leads to MN degeneration. Rescue in either of these pathways would positively impact the others possibly preserving MNs in ALS.

In conclusion, our results suggest a model whereby modulations to actin dynamics can affect the stability and function of the nuclear pore, adding cytoskeletal function to the list of modulators of NCT, together with protein aggregation and oxidative stress ^1,13^. The defects in NCT observed in ALS MNs could be particularly relevant to disease pathogenesis as it may compromise many cellular pathways relying on the integration between signaling and changes in gene expression and/or mRNA processing, such as neurotrophic factors signaling or oxidative stress ^42–44^. Failure to integrate these signaling cascades would decrease the cell’s ability to respond to environmental challenges, leading to the accumulation of toxic insults. Regulating nuclear export or restoring NPC function via modulation of actin could represent feasible therapeutic strategies for multiple forms of ALS and other neurodegenerative diseases beyond ALS.

## MATERIAL and METHODS

### Primary motor neuron culture, transfection, and treatments

Primary motor neurons (MNs) were isolated from E12.5 mouse embryonic spinal cords dissociated in 0.1% trypsin (Worthington) at 37°C for 12 minutes. MNs were purified using a 6% Optiprep (Sigma-Aldrich) density gradient and plated on glass coverslips coated with 0.5g/L poly-ornithine (Sigma-Aldrich) and laminin (Thermo Fisher). Cells were grown at 37°C and 5% CO2 in Neurobasal medium (Thermo Fisher) supplemented with 0.25% Glutamax, 2% B27, 2% horse serum, and 10ng/ml BDNF, GDNF, and CNTF. MNs at 2 days *in vitro* (DIV) were transfected using 1.75µl NeuroMag reagent (OZ Biosciences) + 0.5µg DNA. Complete growth medium was replaced with serum free neurobasal medium 1 hour prior and after transfection. The V5-PFN1 plasmids were described in Wu et al. (2011), while the (G4C2)80 construct was described in ^26^. For the GFP-PFN1 constructs, WT or mutant PFN1 was PCR amplified from the V5 constructs and cloned in the pEGFP-C1 backbone using BamHI sites. The FH1-FH2 domains of murine mDia1 (aa 553-1192) were PCR amplified using specific primers and cloned into the pEGFP-C1 plasmid using XhoI and BamHI restriction sites. GFP or GFP-mDia1 were co-transfected in a 1:2 ratio with PFN1 or *C9ORF72* plasmids. For actin depolymerization assays, 2DIV MNs were treated with 0.1mg/L Latrunculin A (Cayman) for 3 days. For nuclear export inhibition, MNs were treated with 50nM KPT-276 (Selleck) or equal volume of DMSO (Sigma Aldrich) 1 day after transfection.

### Primary cortical neuron culture, transfection, and treatments

Primary cortical neurons were isolated from E15 mouse embryos dissociated in 0.05% trypsin-EDTA (Thermo Scientific) at 37°C for 12 minutes. 320,000 cells/ml were plated on poly-D-lysine (0.125 mg/ml, Sigma Aldrich) and lamin (5 µg/ml, Corning) coated 96-well glass plates and grown at 37°C and 5% CO2 in Neurobasal medium (Thermo Fisher) supplemented with 2% B27 and 1% Glutamax (Thermo Scientific). Four DIV neurons were transfected with 0.2µl Lipofectamine 2000 (Thermo Scientific) and 100ng DNA following manufacturer’s recommendations. A ratio of 4:1 was used for the GFP and S-mCherry vectors. Thirsty-six hours after transfection, Leptomycin B (10mg/L) was added to the culture medium immediately before image acquisition. Cells were imaged using a Nikon TiE widefield microscope equipped with temperature- and CO2-controlled environmental chamber. Movies were acquired with a 20x lens at a rate of 1 frame every 1 or 2 minutes for 1 hour. For rescue experiments, neurons were fixed immediately after acquisition and stained with V5 antibody to detect V5-PFN1 expression.

### Immunofluorescence and image acquisition

Cells were fixed with 4% paraformaldehyde for 15 minutes. Fixed motor neurons were treated with hot 10mM citrate buffer, pH 6 for 20 minutes before permeabilization with 0.2% Triton-X 100 for 5 minutes. Cells were blocked with 5% bovine serum albumin for 45 minutes and hybridized with the appropriate antibodies overnight at 4°C. Anti-mouse and anti-rabbit donkey secondary antibodies or phalloidin conjugated with either Alexa Fluor 647, Alexa Fluor 594, Alexa Fluor 546, or Alexa Fluor 488 (Jackson Immunoresearch and Thermo Fisher) were hybridized for 1 hour at room temperature. Coverslips were mounted onto a glass slide using Prolong Gold mounting medium (Thermo Fisher) and imaged using an epifluorescence microscope (Nikon Ti E) equipped with a cooled CMOS camera (Andor Zyla). Images were acquired as Z-stacks (0.2µm step size) using a 60x lens unless otherwise specified.

### Transmission Electron Microscopy

Six-well plates of cultured cells were fixed overnight at 4°C by adding 1 ml of 2.5% glutaraldehyde in 0.1 M Na cacodylate-HCl buffer (pH 7.2). Fixed samples were washed three times in 0.5 M Na cacodylate-HCl buffer (pH 7.0) and post-fixed for 1 hour in 1% osmium tetroxide (w/v) in the same buffer at room temperature. Following post-fixation, the culture dish with adherent cells were enblock stained (20 min) with 1% aqueous uranyl-acetate (w/v). The fixed cell culture dishes were washed again in the same buffer and dehydrated through a graded series of ethanol to 100% and transferred through two changes of 50/50 (v/v) SPIpon resin (Structure Probe, Inc.) / 100% ethanol and left overnight to infiltrate. The following morning the cell culture plates were transferred through three changes of fresh SPIpon resin to finish the infiltration and embedding and finally, the dishes were filled with freshly prepared SPIpon resin and polymerized for 48 hours at 70°C. Once fully polymerized, the six well-plate was cut apart and each well was plunged into liquid nitrogen to separate the SPIpon epoxy block with the embedded cells from the culture dish. The round epoxy disks with the embedded cells were then examined under an upright light microscope and areas of cells were cut from the disks and glued onto blank microtome studs and trimmed for ultramicrotomy. Ultrathin sections (70 nm) were cut on a Reichart-Jung ultramicrotome using a diamond knife. The sections were collected and mounted on copper support grids and contrasted with lead citrate and uranyl acetate and examined on a FEI Tecnai G2 Spirit transmission electron microscope at 100 Kv accelerating voltage and images were recorded at various magnifications using a Gatan 2 K digital camera system.

### Fluorescence in situ hybridization

Motor neurons were fixed in 4% RNase-free paraformaldehyde for 15 minutes, and stored in 70% ethanol at 4°C overnight. Cells were sequentially incubated for 5 minutes in 1x PBS and wash buffer (2x SSC, 10% formamide), and hybridized in hybridization buffer (10mg/ml dextran sulfate, 4mg/ml BSA, 40µM ribonucleoside vanadyl complexes, 2x SSC, 1% PBS) at 37°C overnight with 12.5µM probes and 0.2mg/ml each salmon sperm DNA and *E. Coli* tRNA. *Neurofilament L* mRNA specific probes were design using Biosearch Technology online tool and conjugated with the Quasar^®^ 570 fluorophore (Table S2). Coverslips were then washed twice for 30 minutes at 37°C in wash buffer before mounting them as described above.

### Axon length and outgrowth analysis

Motor neurons were co-transfected as outlined above with green fluorescent protein (GFP) and V5-tagged PFN1 constructs in a 1:2 ratio. KPT-276 (50nM) or equal volumes of DMSO were added to the culture medium 1 day after transfection and maintained throughout the experiment for a total of 3 days. Cells were fixed and stained to detect V5-PFN1 expression 4 days after transfection. Cells were imaged as individual focal planes using a 10x lens. GFP was used to identify transfected cells and highlight the whole cell structure. For live imaging of axon outgrowth, KPT-276 (50nM) or equal volumes of DMSO were added to the culture medium 1 day after transfection and maintained throughout the experiment for a total of ∼18-24 hours. Cells were imaged at 3 DIV using a Nikon TiE widefield microscope equipped with temperature- and CO2-controlled environmental chamber. Movies were acquired with a 20x lens at a rate of 1 frame every 10 minutes for 1 hour.

### Immunoprecipitations, solubility assays, and western blots

HEK293 cells grown in DMEM + 10% FBS were transfected with V5-PFN1 constructs and lysed with lysis buffer (20mM Tris, 150 mM NaCl, 1% Triton X-100, protease inhibitor Complete EDTA-free, Roche) 24 or 48 hours after transfection. For immunoprecipitations, the detergent-soluble lysates were added to 30 µL of Protein A-Agarose beads (Roche) and 0.3 µg of anti-V5 antibody (Novus) and rocked overnight at 4°C. Immunoprecipitated complexes were eluted in Laemmli buffer (60mM Tris-Cl pH 6.8, 2% SDS, 10% glycerol, 5% beta-mercaptoethanol, 0.01% bromophenol blue) and then subjected to western blot analysis. For solubility assays, lysates were sonicated on ice and then centrifuged at 16,000 *x g* for 10 min at 4°C. The supernatant was collected as the soluble fraction. The pellet was washed three times with lysis buffer, resuspended in 8M urea, sonicated, and then centrifuged at 16,000 *x g* for 10 min at 4°C. The supernatant was collected as the insoluble fraction.

Samples were resolved by SDS-PAGE on Mini Protean TGX 4-20% gradient polyacrylamide gels (Bio-Rad) and transferred onto nitrocellulose membranes (Bio-Rad). Membranes were blocked with Odyssey Blocking Buffer (LI-COR) and probed with primary antibodies overnight. Secondary antibodies conjugated with IRDye^®^ infrared fluorophores (LI-COR) were incubated for 1 hour at room temperature. Blots were visualized using the Odyssey Infrared Imaging System (LI-COR).

### POLDIP3 splicing assay

Immortalized lymphoblastoid cell lines obtained from 3 ALS patients carrying PFN1 mutations and 4 controls ^15^ were cultured in RPMI medium supplemented with 15% fetal bovine serum (FBS). Whole RNA was extracted from 5x10^6^ cells using TriZol reagent (Thermo Fisher) according to manufacturer’s instructions. RNA (2µg) was retrotranscribed using the High-Capacity cDNA Reverse Transcription Kit (Thermo Fisher). RT-PCR was performed using specific primers amplifying exon2-exon4 of human *POLDIP3* mRNA. DNA gels were stained with SYBR Safe dye (Thermo Fisher) and imaged using the ChemiDoc XR+ imager (BioRad).

### Image processing and quantification

Immunofluorescence images were deconvolved using an adaptative blind deconvolution algorithm (Autoquant X3, Media Cybernetics) before analysis. To measure fluorescence intensities, the signals were thresholded and the resulting integrated densities were normalized on the area of the selected region (e.g. cell body, nucleus). Thresholds were kept consistent for all images within experiments. For all experiments, values were normalized to WT averages so that WT always = 1±SEM. For NCT dynamics, the fluorescence intensity of S-mCherry was measured in the nucleus using ImageJ and normalized to the background for every time point. All values were subsequently normalized to T0. For the analysis of axonal fluorescence intensities, a 100µm long region of the proximal axon was measured in all conditions. Axon lengths were measured using the ImageJ plugin NeuronJ ^45^. The axon was defined as the longest neurite. The rate of axon outgrowth was measured using the ImageJ plugin MTrackJ by tracking the movement of the growth cone in all fields ^46,47^. A blinded analysis was performed to assess RanGAP1, RanBP2, POM121, and F-Nups localization. For all experiments, raw values were normalized to the mean of the control condition.

### Statistical Analysis

Statistical analyses were performed using Prism 7 software package (GraphPad). Normality of the samples was assessed using the D’Agostino & Pearson Omnibus test. According to normality, parametric or non-parametric tests were used to assess significance, defined as *p*<0.05. Detailed information on all statistical tests performed is listed in Table S1.

## Data Availability

The datasets generated during the current study are available from the corresponding author on reasonable request.

## ACKNOWLEDGMENTS

The authors would like to thank Thanh Trinh, Anh Nhu, Joseph Lee, and Pamela Keagle for excellent technical support, Dr. Helene Tran for sharing resources, and Dr. Daryl Bosco for insightful discussions and critical feedback. The (G4C2)_80_ expression plasmid was a gift from Dr. Charlet-Berguerand. NLS-mCherry-NES reporter (pDN160) was a gift from Dr. Di Ventura & Dr. Eils (Addgene plasmid # 72660). This work was supported by the American ALS Association (15-PDF-211 and 18-IIA-418 to CF), National Institute of Health/NINDS (F32 NS090831 to EWD and R01 NS073873 to JEL) and National Institute of Health/NCRR (S10RR027897 and S10OD021580) to the UMass Med EM core. VG was supported by the Doctorate School of Molecular and Translational Medicine, Università degli Studi di Milano.

## AUTHOR CONTRIBUTIONS

EWD, AG, MJ, VG, SB, and CF performed the experiments and analyzed results. RHB, JEL, and CF provided resources and helped with data interpretation. CF and JEL designed and directed the project and secured funding. CF wrote the manuscript. All authors reviewed the final manuscript.

## REFERENCES

1. Chou, C. C. et al. TDP-43 pathology disrupts nuclear pore complexes and nucleocytoplasmic transport in ALS/FTD. Nat Neurosci 21, 228–239 (2018).

2. Freibaum, B. D. et al. GGGGCC repeat expansion in C9orf72 compromises nucleocytoplasmic transport. Nature 525, 129–133 (2015).

3. Jovičić, A. et al. Modifiers of C9orf72 dipeptide repeat toxicity connect nucleocytoplasmic transport defects to FTD/ALS. Nat Neurosci 18, 1226–1229 (2015).

4. Kinoshita, Y. et al. Nuclear contour irregularity and abnormal transporter protein distribution in anterior horn cells in amyotrophic lateral sclerosis. J Neuropathol Exp Neurol 68, 1184–1192 (2009).

5. Zhang, J. et al. Altered distributions of nucleocytoplasmic transport-related proteins in the spinal cord of a mouse model of amyotrophic lateral sclerosis. Acta Neuropathol 112, 673–680 (2006).

6. Zhang, K. et al. The C9orf72 repeat expansion disrupts nucleocytoplasmic transport. Nature 525, 56–61 (2015).

7. Polymenidou, M. et al. Misregulated RNA processing in amyotrophic lateral sclerosis. Brain Res 1462, 3–15 (2012).

8. Cronshaw, J. M. & Matunis, M. J. The nuclear pore complex: disease associations and functional correlations. Trends Endocrinol Metab 15, 34–39 (2004).

9. Jamali, T., Jamali, Y., Mehrbod, M. & Mofrad, M. R. Nuclear pore complex: biochemistry and biophysics of nucleocytoplasmic transport in health and disease. Int Rev Cell Mol Biol 287, 233–286 (2011).

10. D’Angelo, M. A., Raices, M., Panowski, S. H. & Hetzer, M. W. Age-dependent deterioration of nuclear pore complexes causes a loss of nuclear integrity in postmitotic cells. Cell 136, 284–295 (2009).

11. Savas, J. N., Toyama, B. H., Xu, T., Yates, J. R. & Hetzer, M. W. Extremely long-lived nuclear pore proteins in the rat brain. Science 335, 942 (2012).

12. Woerner, A. C. et al. Cytoplasmic protein aggregates interfere with nucleocytoplasmic transport of protein and RNA. Science 351, 173–176 (2016).

13. Zhang, K. et al. Stress granule assembly disrupts nucleocytoplasmic transport. Cell 173, 958–971.e17 (2018).

14. Boopathy, S. et al. Structural basis for mutation-induced destabilization of profilin 1 in ALS. Proc Natl Acad Sci U S A 112, 7984–7989 (2015).

15. Wu, C. H. et al. Mutations in the profilin 1 gene cause familial amyotrophic lateral sclerosis. Nature 488, 499–503 (2012).

16. Mitchell, J. M., Mansfeld, J., Capitanio, J., Kutay, U. & Wozniak, R. W. Pom121 links two essential subcomplexes of the nuclear pore complex core to the membrane. J Cell Biol 191, 505–521 (2010).

17. Onischenko, E. et al. Natively unfolded FG repeats stabilize the structure of the nuclear pore complex. Cell 171, 904–917.e19 (2017).

18. Cho, K. I. et al. Loss of Ranbp2 in motoneurons causes disruption of nucleocytoplasmic and chemokine signaling, proteostasis of hnRNPH3 and Mmp28, and development of amyotrophic lateral sclerosis-like syndromes. Dis Model Mech 10, 559–579 (2017).

19. Mahajan, R., Gerace, L. & Melchior, F. Molecular characterization of the SUMO-1 modification of RanGAP1 and its role in nuclear envelope association. J Cell Biol 140, 259–270 (1998).

20. Shani, T., Levy, M. & Israelson, A. Assay to Measure Nucleocytoplasmic Transport in Real Time within Motor Neuron-like NSC-34 Cells. J Vis Exp (2017). doi:10.3791/55676

21. Alami, N. H. et al. Axonal transport of TDP-43 mRNA granules is impaired by ALS-causing mutations. Neuron 81, 536–543 (2014).

22. Shiga, A. et al. Alteration of POLDIP3 splicing associated with loss of function of TDP-43 in tissues affected with ALS. PLoS ONE 7, e43120 (2012).

23. Archbold, H. C. et al. TDP43 nuclear export and neurodegeneration in models of amyotrophic lateral sclerosis and frontotemporal dementia. Sci Rep 8, 4606 (2018).

24. Haines, J. D. et al. Nuclear export inhibitors avert progression in preclinical models of inflammatory demyelination. Nat Neurosci 18, 511–520 (2015).

25. Kovar, D. R., Harris, E. S., Mahaffy, R., Higgs, H. N. & Pollard, T. D. Control of the assembly of ATP- and ADP-actin by formins and profilin. Cell 124, 423–435 (2006).

26. Almeida, S. et al. Modeling key pathological features of frontotemporal dementia with C9ORF72 repeat expansion in iPSC-derived human neurons. Acta Neuropathol 126, 385–399 (2013).

27. Sivadasan, R. et al. C9ORF72 interaction with cofilin modulates actin dynamics in motor neurons. Nat Neurosci 19, 1610–1618 (2016).

28. Taylor, J. P., Brown, R. H. & Cleveland, D. W. Decoding ALS: from genes to mechanism. Nature 539, 197–206 (2016).

29. Boeynaems, S., Bogaert, E., Van Damme, P. & Van Den Bosch, L. Inside out: the role of nucleocytoplasmic transport in ALS and FTLD. Acta Neuropathol 132, 159–173 (2016).

30. Minakhina, S., Myers, R., Druzhinina, M. & Steward, R. Crosstalk between the actin cytoskeleton and Ran-mediated nuclear transport. BMC Cell Biol 6, 32 (2005).

31. Tanaka, Y., Nonaka, T., Suzuki, G., Kametani, F. & Hasegawa, M. Gain-of-function profilin 1 mutations linked to familial amyotrophic lateral sclerosis cause seed-dependent intracellular TDP-43 aggregation. Hum Mol Genet 25, 1420–1433 (2016).

32. Figley, M. D., Bieri, G., Kolaitis, R. M., Taylor, J. P. & Gitler, A. D. Profilin 1 associates with stress granules and ALS-linked mutations alter stress granule dynamics. J Neurosci 34, 8083–8097 (2014).

33. Jahed, Z., Soheilypour, M., Peyro, M. & Mofrad, M. R. The LINC and NPC relationship - it’s complicated J Cell Sci 129, 3219–3229 (2016).

34. Wang, J. Y. et al. Sun1 deficiency leads to cerebellar ataxia in mice. Dis Model Mech 8, 957–967 (2015).

35. Gros-Louis, F. et al. Mutations in SYNE1 lead to a newly discovered form of autosomal recessive cerebellar ataxia. Nat Genet 39, 80–85 (2007).

36. Kumar, J. P., Wilkie, G. S., Tekotte, H., Moses, K. & Davis, I. Perturbing nuclear transport in Drosophila eye imaginal discs causes specific cell adhesion and axon guidance defects. Dev Biol 240, 315–325 (2001).

37. Dent, E. W., Gupton, S. L. & Gertler, F. B. The growth cone cytoskeleton in axon outgrowth and guidance. Cold Spring Harb Perspect Biol 3, (2011).

38. Jung, H., Yoon, B. C. & Holt, C. E. Axonal mRNA localization and local protein synthesis in nervous system assembly, maintenance and repair. Nat Rev Neurosci 13, 308–324 (2012).

39. Jin, L. Q., Pennise, C. R., Rodemer, W., Jahn, K. S. & Selzer, M. E. Protein synthetic machinery and mRNA in regenerating tips of spinal cord axons in lamprey. J Comp Neurol 524, 3614–3640 (2016).

40. Wang, E. T. et al. Dysregulation of mRNA Localization and Translation in Genetic Disease. J Neurosci 36, 11418–11426 (2016).

41. Fallini, C., Donlin-Asp, P. G., Rouanet, J. P., Bassell, G. J. & Rossoll, W. Deficiency of the Survival of Motor Neuron Protein Impairs mRNA Localization and Local Translation in the Growth Cone of Motor Neurons. J Neurosci 36, 3811–3820 (2016).

42. Aulas, A. & Vande Velde, C. Alterations in stress granule dynamics driven by TDP-43 and FUS: a link to pathological inclusions in ALS? Front Cell Neurosci 9, 423 (2015).

43. Sloan, K. E., Gleizes, P. E. & Bohnsack, M. T. Nucleocytoplasmic Transport of RNAs and RNA-Protein Complexes. J Mol Biol 428, 2040–2059 (2016).

44. Hill, S. J. et al. Two familial ALS proteins function in prevention/repair of transcription-associated DNA damage. Proc Natl Acad Sci U S A 113, E7701–E7709 (2016).

45. Meijering, E. et al. Design and validation of a tool for neurite tracing and analysis in fluorescence microscopy images. Cytometry A 58, 167–176 (2004).

46. Meijering, E., Dzyubachyk, O. & Smal, I. Methods for cell and particle tracking. Meth Enzymol 504, 183–200 (2012).

47. Schneider, C. A., Rasband, W. S. & Eliceiri, K. W. NIH Image to ImageJ: 25 years of image analysis. Nat Methods 9, 671–675 (2012).

